# Thalamocortical network dynamics in focal epilepsy: SEEG investigation

**DOI:** 10.64898/2026.03.05.709626

**Authors:** Elliot M. Nester, Maya A. Jayaram, Tejas Umesh, Lekha Varisa, Kris Phataraphruk Rains, Kris A. Smith, Kevin Choi, Deana M. Gazzola, Susan T. Herman, Laura Lehnhoff, Courtney Schusse, Vladimir Shvarts, Ritika Suri, Yalin Wang, Bradley Greger, Yuan Wang, Pavan Turaga, Stephen T. Foldes, David P. Harris, T. Noah Hutson, Andrew I. Yang

## Abstract

Thalamic neuromodulation is clinically effective in drug­resistant epilepsy, suggesting critical contributions of the thalamus to the epileptogenic process. However, the underlying electrophysiologic mechanisms remain poorly characterized. Converging evidence implicates the thalamus in shaping large­scale functional interactions across the cortex. We hypothesized that ictal changes in thalamic activity track cortical network dynamics associated with seizure propagation.

We analyzed stereo­electroencephalography recordings from 16 patients with focal epilepsy (255 seizures) with simultaneous sampling of the thalamus (anterior nucleus, N=14; pulvinar, N=11) and cortex. Cortical regions of interest included the seizure onset zone (SOZ), surrounding cortices (near­SOZ), and control regions from the contralateral hemisphere. We characterized seizure dynamics across spatial scales, from local activity within each region to network­level, inter­regional interactions. Local activity was decomposed into its periodic (oscillatory) and aperiodic components. Network interactions were characterized by directed functional connectivity computed with a multivariate method.

Seizures were associated with increased broadband power (a proxy for neuronal population firing rates) and low­frequency rhythmic activity across the thalamocortical network relative to interictal baseline levels. In contrast, consistent changes in aperiodic slope (a putative marker of excitation­inhibition balance) were specific to the thalamus, which showed an early and sustained steepening (i.e., more negative slope). While local rhythms were heterogeneous across the canonical frequency bands, inter­regional interactions predominantly involved the beta band (13­30 Hz). Shortly after onset, both forward outflow from SOZ to near­SOZ and feedback inflow in the reverse direction were increased. These bidirectional effects were expressed via both a direct cortico­cortical pathway and an indirect transthalamic route, operating in parallel. These dynamics were further stratified based on seizure subtypes, leveraging the fact that there was minimal propagation of ictal activity to the near­SOZ in subclinical seizures. The ictal drop in thalamic aperiodic slope was primarily observed in clinical seizures. At the network level, whereas SOZ→near­SOZ outflow was present across seizure types, reverse feedback was particularly enhanced in clinical seizures. Multivariable regression showed that the degree of thalamic slope steepening uniquely tracked seizure­to­seizure fluctuations in the strength of near­ SOZ→SOZ feedback, and further predicted seizure durations.

Together these findings highlight thalamic aperiodic slope as an index of cortical network dynamics linked to seizure propagation, with potential clinical utility for further development of physiology­informed precision neuromodulation.

## Introduction

Thalamic neuromodulation has transformed the surgical management of patients with drug­ resistant epilepsy. Since its regulatory approval, Deep Brain Stimulation (DBS) of the anterior nucleus of the thalamus (ANT) has been increasingly utilized for focal epilepsy.^1–3^ However, there is substantial variability in seizure outcomes across patients, motivating a resurgence of clinical investigations of other thalamic nuclei, e.g. centromedian (CM) nucleus and pulvinar.^4–6^ These developments present opportunities for precision neuromodulation in epilepsy. Concomitantly, there has been growing interest in personalized target selection using patient­ specific data, such as electrophysiologic recordings obtained simultaneously from the thalamus and cortical regions with stereo­electroencephalography (SEEG). A budding literature has focused on visual review of ictal EEG to assess for the presence and latency of propagated ictal activity across thalamic subdivisions.^7–9^ However, whether stronger and quicker spread to a given thalamic nucleus can guide personalized neuromodulation in individual patients remains unknown. In fact, while clinical experience in thalamic SEEG extends back two decades,^10,11^ further studies to elucidate the contributions of the thalamus to the epileptogenic process will inform clinical trials evaluating the benefits of thalamic SEEG and ultimately more widespread clinical adoption.

There is growing recognition that the anatomic distribution of pathological changes in epilepsy is diffuse, extending beyond the brain regions where seizures start, referred to as the seizure onset zone (SOZ).^12–14^ Under this framework, the pathologic substrate of seizures is postulated to be a network comprised of multiple brain regions that participates in seizure initiation, propagation, and termination ­ denoted as the epileptogenic network.^15–17^ Whereas mounting evidence suggests the critical contributions of cortical regions outside the SOZ (non­ SOZ),^18–22^ the role of the thalamus remains relatively understudied, in part due to the relative scarcity of invasive recordings from the human thalamus. In generalized epilepsy, animal models of absence seizures suggest a central role of the thalamus in ictogenesis.^23,24^ In focal seizures, human SEEG studies have suggested that the thalamus may be involved in seizure termination.^25–28^ However, invasive recordings of the ictal thalamocortical network in humans remain scarce,^25–35^ and little is known about thalamic contributions to the earlier stages of seizures.

Using thalamic recordings obtained from deep brain stimulation (DBS) devices, we previously showed that seizures were associated with robust changes in aperiodic slope^36^ – a putative marker of excitation­inhibition (E­I) balance.^37–39^ More broadly, a growing literature supports a critical role of the thalamus in shaping cortex­wide neural dynamics, both in health and in disease.40,41 The specific thalamic nuclei that have been implicated are those with widespread bidirectional connectivity with cortical and limbic structures, which overlaps with the most commonly­utilized neuromodulation targets. Biologically­plausible models of thalamocortical networks have shown that the thalamus can act as a switchboard to regulate the flow of neural activity in the cortex.^42^ The key variable that functioned as a control knob to bias inter­regional interactions between cortical regions was the gain or “excitability” of thalamic neurons (of which E­I balance is a major determinant of).^43,44^ Converging evidence from human fMRI studies has shown that measures of neural variability^45^ (of which aperiodic slope is one) in the thalamus are robust markers of brain­wide functional interactions, both within and across individuals.^46,47^ Motivated by these findings, we hypothesized that thalamic aperiodic slope tracks network patterns associated with the cortical propagation of seizures.

To test our hypothesis, we studied local field potentials (LFP) from focal epilepsy patients with simultaneous recordings from the cortex and thalamus (ANT and pulvinar). Analysis of cortical dynamics were further stratified across the SOZ, regions immediately outside the SOZ (near­SOZ), and control regions in the hemisphere contralateral to the SOZ (far­ SOZ). These cortical regions exhibited differential participation in ictal network dynamics, which were further distinguished across seizure types based on semiology. We first characterized ictal changes in the local activity of each region, stratifying across broadband activity (a marker of neuronal firing^48^), low­frequency oscillatory activity, and aperiodic slope dynamics. We then assessed inter­regional interactions throughout the thalamocortical ictal network with a multivariate measure of directed functional connectivity. Third and finally, we bridged across these spatial scales by evaluating the relationship between thalamic local activity and cortical network patterns across individual seizures as well as across seizure types.

## Materials and methods

### Study cohort

Subjects were patients with drug­resistant epilepsy who were deemed to be candidates for invasive monitoring in the multidisciplinary epilepsy surgery conference held at the Level 4 Comprehensive Epilepsy Center of the Barrow Neurological Institute. Neural data were obtained while admitted to the Epilepsy Monitoring Unit after surgical implantation of SEEG electrodes (12/2023­9/2024). Inclusion criteria included: voluntary informed consent provided for research use of data in accordance with the Declaration of Helsinki; sampling of ANT and/or pulvinar with SEEG; ≥4 recorded seizures. All research procedures were approved by the local Institutional Review Board (IRB). Research participation did not impact clinical care, including SEEG implant number or locations. At our center, thalamic sampling is prescribed on a case­by­ case basis during the epilepsy surgery conference, and is driven strictly by clinical indications only.

Our study population included 16 patients (Table 1): 11 had unifocal and 5 had bifocal epilepsy, for a total of 21 seizure onset zones (SOZ), the majority of which were localized to the temporal lobe (12 mesial temporal, 2 neocortical, 4 both). Remaining SOZs were in insular (N=1) or basal temporo­occipital cortices (N=2). Structural lesions underlying epilepsy included mesial temporal sclerosis (MTS; N=3), hippocampal abnormality not meeting criteria for MTS (N=3), and focal cortical dysplasia (FCD; N=1). A total of 255 seizures were included for analysis (N=15.9±15.6 [mean±s.d.] per subject; duration 69.3±41.2 s). Seizure types for subgroup analyses were determined from the epileptologist’s clinical report, and included subclinical (electrographic) seizures, clinical seizures with preserved consciousness (focal aware [FA]), and clinical seizures with impaired consciousness (both focal impaired awareness [FIA] and focal to bilateral tonic clonic [FBTC]).

**Table 1:**
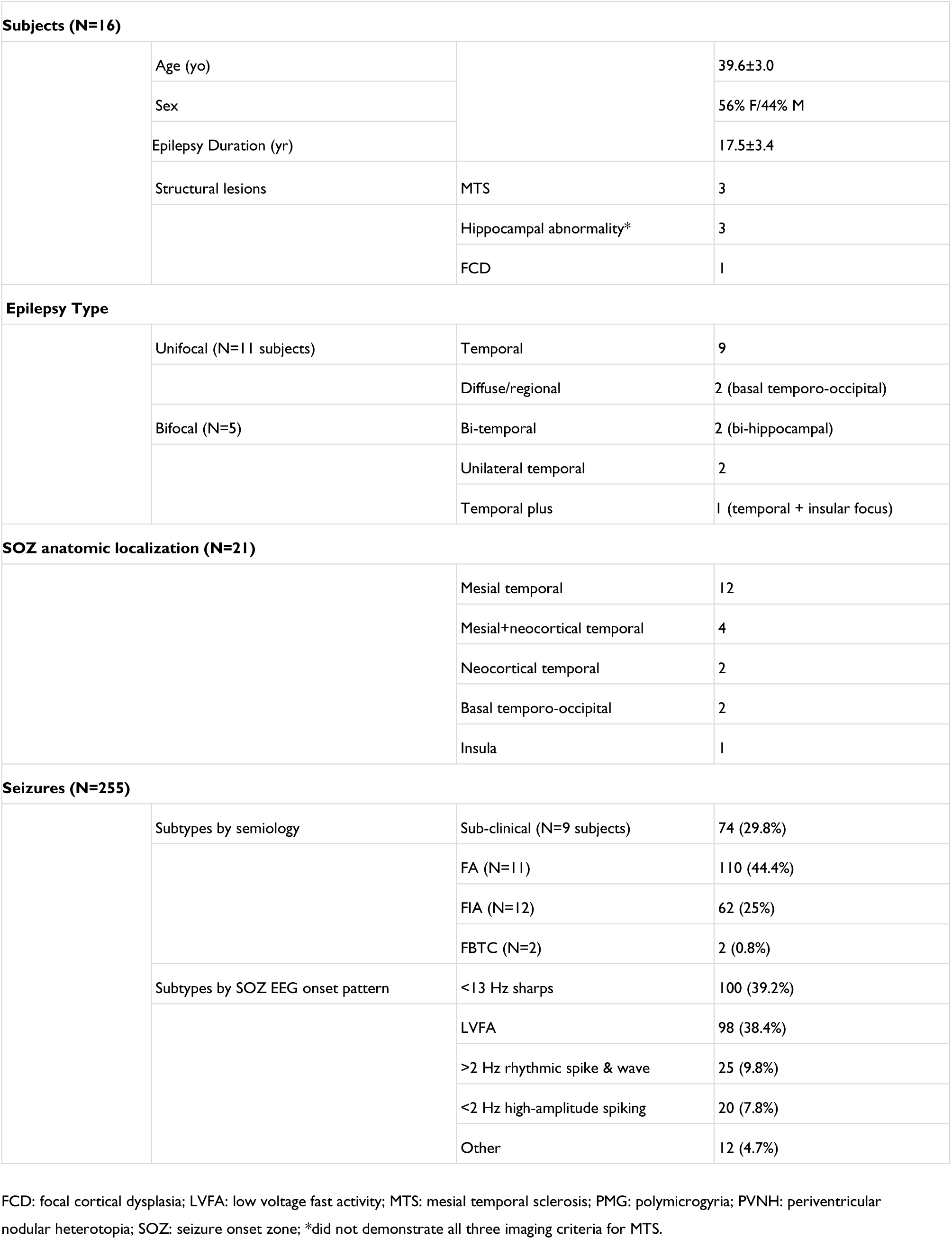
Demographic and clinical characteristics.

### Data acquisition and anatomic localization

LFP recordings were obtained with depth electrodes (Ad­Tech) containing 6­16 platinum macrocontacts (2.41 mm length, 3­5 mm spacing). All data were recorded at 1 kHz (Nihon Kohden). Data were notch­filtered at 60 Hz (2^nd^ order IIR filter) and high­pass filtered at 0.1 Hz (2^nd^ order Butterworth filter). Bipolar referencing was then performed across adjacent contacts within the same electrode to yield “virtual” contacts.

Anatomic localization of electrode contacts was performed following established methods, as presented here in brief. Each contact was segmented on the post­operative CT, and co­registration was performed with the pre­operative MRI (Curry, Compumedics Neuroscan). Parcellation of cortical regions and segmentation of subcortical structures, including thalamic nuclei, was performed on Freesurfer^49^ based on widely­used atlases.^50–52^ Anatomic localization of each contact was performed in each subject’s native MRI space. As analyzed signals were bipolar referenced, recordings locations were treated as the midpoint of the corresponding pair of contacts. Anatomic localization of thalamic virtual contacts were visually confirmed on the subject’s MRI. For group­level visualizations and characterizations of recording locations, Montreal Neurological Institute (MNI) coordinates were generated for each virtual contact after non­linear registration to the MNI 305 brain.^53^

### Regions of interest for analysis

We defined four regions of interest (ROI), independently for each seizure (Fig. 1A): (i) SOZ; (ii) cortical regions immediately surrounding the SOZ (near­SOZ); (iii) cortical regions far from the SOZ (far­SOZ); (iv) and ipsilateral thalamus. SOZ included data from all contacts noted to have first ictal change in the epileptologist’s clinical report (5.3±5.5 contacts per subject). For each seizure, we chose a single virtual contact as the near­SOZ channel. To do so, we took all virtual contacts not included in the SOZ, and chose the one with the minimum Euclidean distance from the geometric centroid of the corresponding SOZ. These were obtained from the hemisphere ipsilateral to the SOZ in all cases. The average Euclidean distance of SOZ virtual contacts was 5.9±5.7 cm from its centroid, and 14.5±6.1 cm for near­SOZ virtual contacts. For each seizure, we also selected one far­SOZ channel, taking the virtual contact with the greatest distance from the SOZ centroid. Far­SOZ channels were from the contralateral hemisphere in all cases, none which were from brain regions homologous to those comprising the SOZ. Finally, thalamic data were obtained from all virtual contacts within the ipsilateral ANT (2.2±1.0 contacts) and/or pulvinar nucleus (2.3±0.86 contacts). Thalamic recordings were obtained from ANT (N=14) and/or pulvinar (N=11; both, N=10; Fig. 1B).

**Figure 1:**
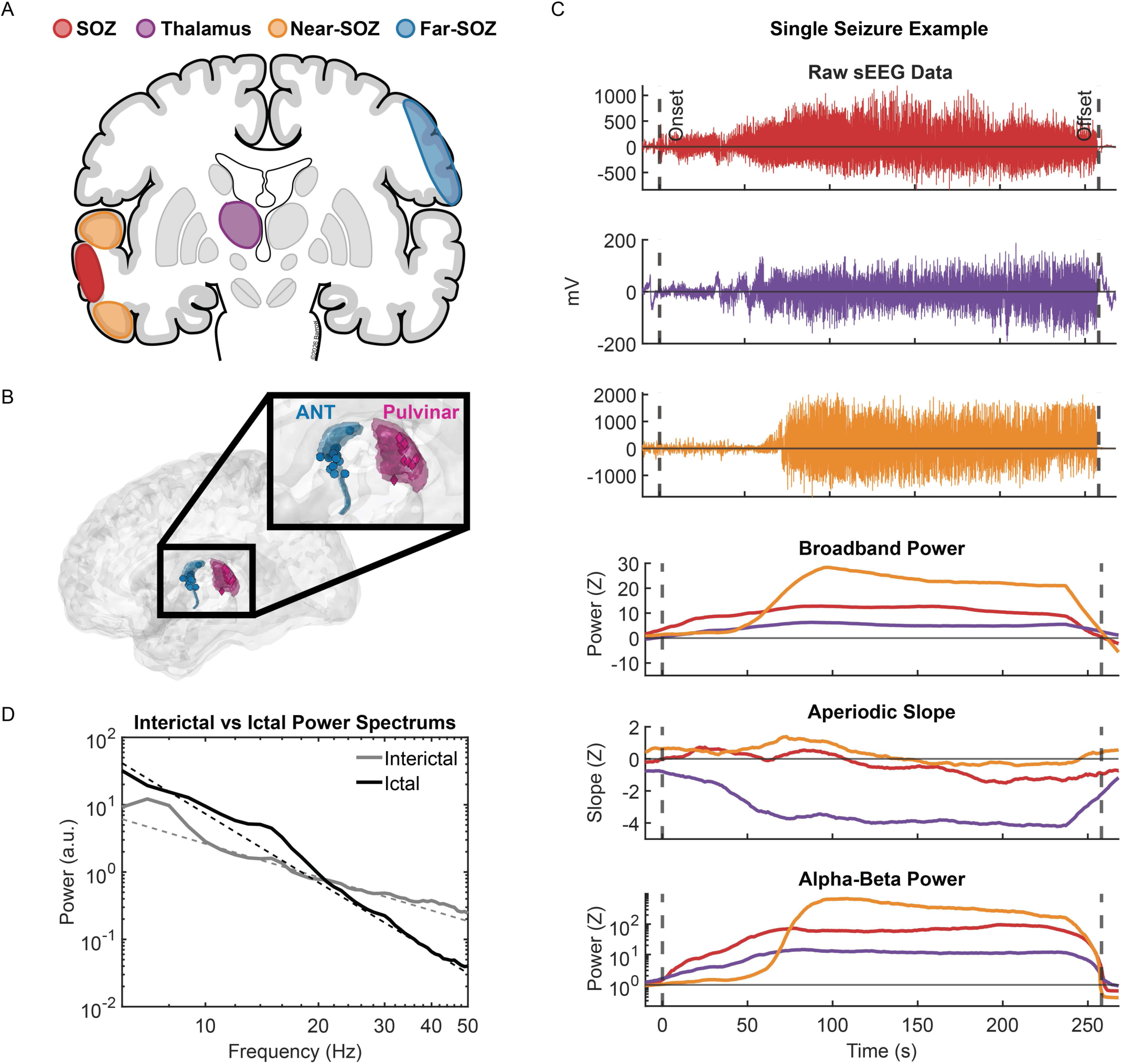
Thalamocortical network schematic and example seizure-level dynamics. **(A)** Regions of interest (ROI). **(B)** Thalamic recording locations showing average coordinates for each subject on MNI brain. **(C)** Single seizure example of ictal dynamics showing raw LFP traces from SOZ, thalamus and near-SOZ (top three panels). Bottom three panels show the three distinct neural activity metrics computed in each ROI. **(D)** Example thalamic LFP power spectrum showing decreased (steeper) aperiodic slope during seizure vs. interictal baseline. Aperiodic slope is obtained from a linear fit of the aperiodic component (dashed lines) in log-log space, after removal of any oscillatory peaks.^54^

### Metrics of local (intra-regional) neural activity

Broadband power and aperiodic slope were quantified from the power spectral density (PSD) following established methods.^48,54^ PSD was computed with the multitaper method (discrete prolate spheroidal sequence tapers) applied across 1 s windows (0% overlap) with the following parameters: 145 linearly­spaced frequency bands spanning 6 to 150 Hz (4 Hz bandwidth); 500 ms windows (50% overlap), corresponding to three cycles at 6 Hz. To extract broadband power, a linear fit of the PSD was performed in log­log space, taking the midpoint of the resulting fit across the entire frequency range.^48^ The aperiodic (1/*f*) slope refers to the power law exponent of the PSD, which was estimated by the slope of a linear fit performed in log­log space. Specifically, we computed the aperiodic slope using the FOOOF algorithm, which disentangles periodic and aperiodic components of the PSD using an iterative approach.^54^ The aperiodic component is the residual of the PSD after removal of any oscillatory peaks. The linear fit was performed across a narrower frequency range of 6­50 Hz. Finally, to assess narrowband rhythmic activity, spectral power was quantified using the Hilbert transform, which was applied to bandpass­filtered (FIR) signals across 36 logarithmically­spaced sub­bands spanning 1 to 150 Hz (bandwidths with 50% overlap). Example neural dynamics during an individual seizure is shown in Fig. 1C, D. As there were potentially multiple SOZ and thalamic channels for each seizure, subject­level results were obtained by averaging across virtual contacts using a resampling approach (see **Statistical analysis**).

### Metrics of inter-regional directional functional connectivity

We characterized inter­regional interactions using a frequency­resolved metric of directed functional connectivity: generalized partial directed coherence (GPDC).^55^ GPDC is a member of a class of methods that uses the ability to predict a time series signal with the past of another time series signal as a proxy measure of “causal” influence (i.e., Granger causality), and is based on fitting a multivariate autoregressive (MVAR) model. Our choice of GPDC was driven by the following key advantages. First, to infer “causal” (Granger) relationships with *frequency specificity* (contrast with refs^25,27–29^). Second, GPDC is a multivariate method controlling for confounds related to common inputs (hence, partial coherence), by disambiguating true (direct) time­lagged correlations from those relayed by a third source (contrast with refs^25–29,32^). Third, GPDC is more robust to cross­channel differences in prediction error (residual) variance, which is the portion of each region’s activity not explained by the past activity of the modeled network. This is made possible by explicitly incorporating the MVAR innovation covariance in the connectivity measure; this innovation­weighted normalization distinguishes GPDC from standard PDC. For each seizure, the MVAR model was fitted with data from each of the four ROIs, independently across 1 s windows (0% overlap). GPDC was then computed from the model coefficients across 1 to 40 Hz (1 Hz steps). Again, as there were potentially multiple SOZ and thalamic recording channels for each seizure, subject­level results were obtained by averaging across models constructed independently for each unique pair of SOZ and thalamic virtual contact using a resampling approach (see **Statistical analysis**).

### Mixed-effects modeling

We employed mixed­effects models to assess temporal trends in beta­band GPDC strength across seizure epochs, comparing the first vs. last 20%, and the first vs. second 50%. Models were fit separately for each GPDC region pair and directionality, as follows:

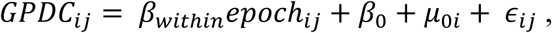

where *epoch_ij_* is a numerical variable (1 or 2) indicating the seizure epoch for subject *i*, seizure *j*; β_0_ is the group-level (fixed) intercept; μ_0*i*_ is the subject-level (random) intercept; and ε_*i*j_ is the residual error. Overall significance of the model was determined by performing a likelihood ratio test against a null model that contained only the intercept terms. To assess whether GPDC strength was increased in the later seizure epoch, we tested the statistical significance of β_*within*_ using Wald’s *t*-test.

To investigate seizure-to-seizure relationships between thalamic aperiodic slope and other neural metrics (local rhythmic power, GPDC), or between seizure duration and thalamic slope, we employed mixed-effects models with both within- and between-subject terms:

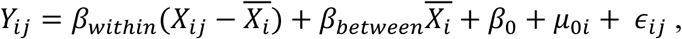

where *Y*_*i*j_ and *X*_*i*j_ are the dependent and independent variables, respectively, for subject *i*, seizure *j*; 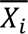 is seizure-averaged independent variable for subject *i*; β_*within*_ is the slope for within-subject, between-seizure differences; β__*between*__ is the slope for between-subject differences. Likelihood ratio test was performed against a null model that only included the between-subject and intercept terms. To assess for between-seizure relationships that were consistently observed across subjects, we again tested the significance of β_*within*_ using Wald’s *t*-test.

### Statistical analysis

To assess ictal dynamics that were consistently observed across subjects, all group-level results were obtained with each subject contributing one data point, averaging across repeated measurements (i.e., seizures×recordings channels). This stands in contrast to statistical tests performed at the seizure level (see refs^25,26,29–33^), which can artificially boost statistical power by falsely assuming that multiple seizures from individual subjects are independent observations. To control for differences in the number of virtual contacts within a ROI across seizures and subjects, as well as in the number of seizures across subjects, subject and group-level results were rigorously computed using a resampling approach (50 iterations).

Prior to inclusion in subject and group-level analyses, seizure-level neural metrics were normalized with respect to baseline data obtained from the inter-ictal period (≥4 hrs from the closest seizures), separately for each bipolar-referenced channel to control for differences across recording contacts (e.g., tissue impedance). Normalization was performed by *z*-scoring ictal values with respect to baseline values. This was the case for all metrics except narrowband activity assessed across 36 sub-bands spanning 1 to 150 Hz, for which spectral power was normalized using percent change 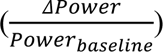 given low variability in baseline activity in a subset of the sub-bands. Baseline-normalized ictal data points that were >5 s.d. were considered artifactual and omitted from subject and group-level analysis.

For subject-level statistical tests, we employed a non-parametric surrogate data (permutation-based) approach, whereby test-specific null distributions were constructed by randomly shuffling ictal vs. interictal labels (N=1000). Interictal data were again obtained by randomly sampling segments from the interictal period with durations equivalent to each seizure. Correction for multiple comparisons across time and frequency was performed with cluster-level statistics.^56^ Clusters were identified after applying pixel-level thresholds designed to ensure comparable sensitivity for detecting one-dimensional (temporal) clusters for broadband power and aperiodic slope, and two-dimensional (time-frequency [TF]) clusters for spectral power and GPDC strength: bottom/top 33^rd^ percentile for the former, and 10^th^ percentile for the latter. These thresholds yield similar percentages of suprathreshold samples for normally-distributed data. Final subject-level results were obtained after *z-*score normalizing the observed statistics with respect to the corresponding null distributions.

To quantify ictal changes in spectral power and GPDC strength in analyses constrained to frequency ranges of interest (8-30 Hz for spectral power, 13-30 Hz for GPDC), we computed averaged values across all time points during seizures as well as across all frequency sub-bands spanning the frequency ranges of interest. This allowed including data from subjects who did not have a significant TF cluster in group-level results. The center frequencies of TF clusters were determined by taking the frequency sub-band with the maximum time-averaged value. To assess group-level temporal dynamics, we controlled for variable seizure durations by linearly interpolating the values of seizure-level neural metrics across a normalized time axis spanning the ictal period from 0% to 100%, corresponding to seizure onset and offset, respectively. Onset latencies for each subject were reported as the first time point at which baseline-normalized ictal values (averaged across frequency sub-bands spanning the ranges of interest) exceeded 2 *z*.

Group-level tests utilized the appropriate non-parametric test, as noted in the **Results**. Correction for multiple comparisons was performed using the false discovery rate (FDR) procedure applied to each family of tests. Reported significant results survived FDR correction (*q*<0.05). Statistical significance was determined using a threshold of *P*<0.05 for two-tailed tests, which was corrected for one-tailed tests.

## Results

### Ictal rise in broadband power is regionally diffuse, aperiodic slope steepening is thalamus specific

Seizures were associated with increased broadband (6-150Hz) power (a marker of neuronal population firing^48^) relative to the interictal baseline in all ROIs (Fig. 2A). This was observed both at the group level (*z*>3.46, *P*<10^-33^, one-sample Wilcoxon signed-rank test), and consistently across subjects (*P*<0.05, permutation test, cluster-corrected; solid circles). Across subjects, broadband power increased two standard deviations above baseline levels (>2 *z*) in early seizure: 14±4% (mean±s.e.m. across subjects) in SOZ; 14±5% in thalamus; and 20±4% in near-SOZ (vertical lines, Fig. 2B). Note that broadband activity was increased in SOZ and thalamus at similar latencies.

**Figure 2:**
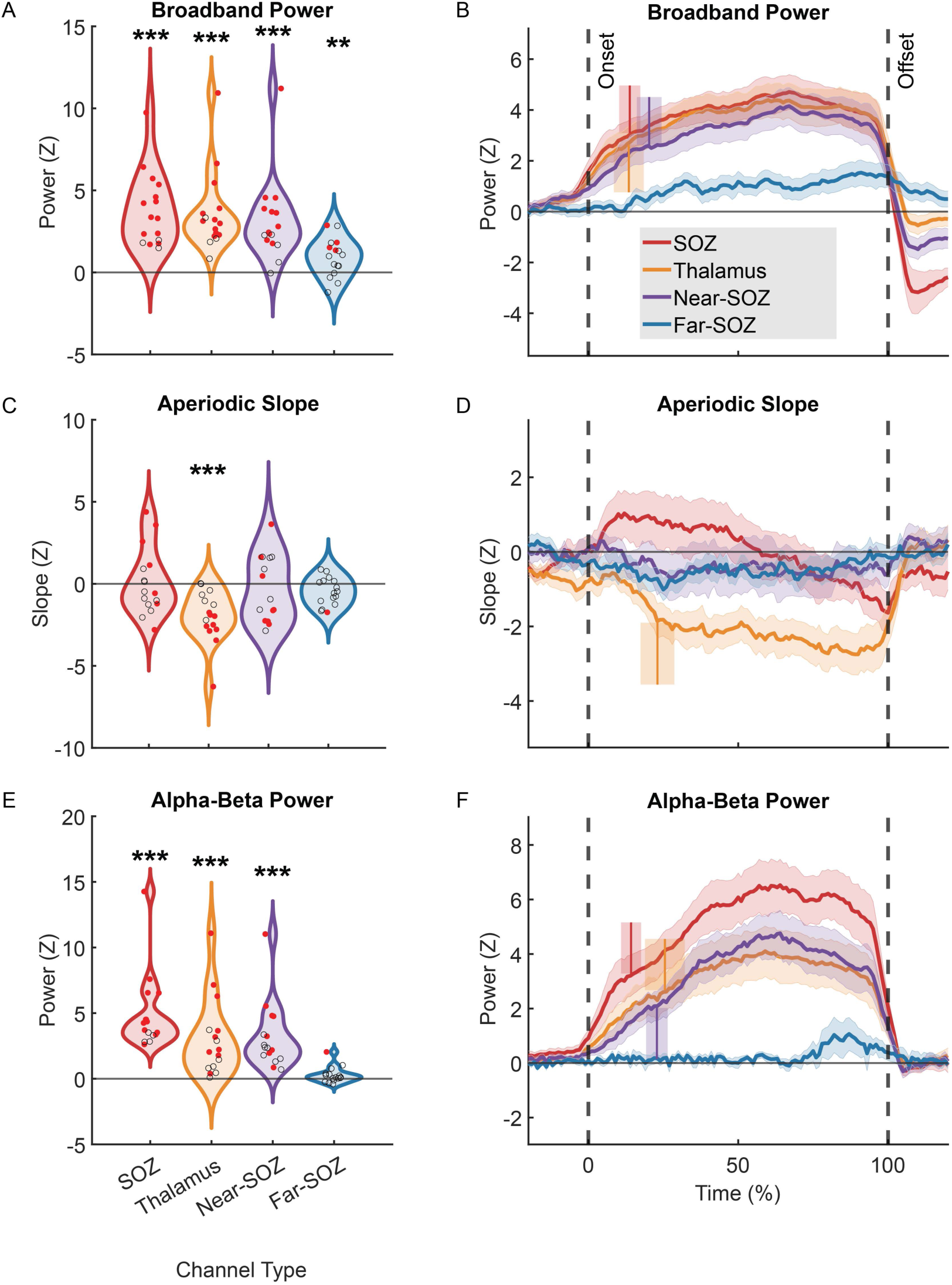
Local ictal dynamics stratified across broadband activity, low-frequency rhythms, and aperiodic slope. **(A)** Time-averaged ictal broadband power (z-scored w.r.t. interictal baseline) across all subjects. Each data point represents subject-level results averaging across seizures, with significance on subject-level tests marked in solid red (P<0.05, permutation test, cluster-corrected). Group-level significance assessed with one-sample Wilcoxon signed-rank test, FDR-corrected across ROIs. **(B)** Subject-averaged temporal dynamics of broadband power. Vertical bars mark average onset latency (>2 z w.r.t. baseline); shadings show s.e.m. across subjects. **(C, D)** As in **(A, B)**, for aperiodic slope. Consistent changes across subjects were only observed in the thalamus. **(E, F)** As in **(A, B)**, for alpha-beta power. All regions showed significant group-level changes except the far-SOZ. w.r.t.: with respect to; \**P*<0.05, \*\**P*<0.01, \*\*\**P*<10^-3^. See Supplementary Fig. 1A for results stratified across distinct thalamic nuclei.

In contrast, consistent changes in aperiodic slope (a marker of E-I balance^37–39^) across subjects were regionally specific to the thalamus (Fig. 2C). Seizures were associated with a sustained decrease in aperiodic slope (group-level *z*=3.41, *P*<10^-33^; 9/16 subjects showed statistical significance) that began in early seizure (23±6%; Fig. 2D). On subgroup analysis across the two thalamic nuclei, the degree of ictal steepening of aperiodic slope was not statistically distinguishable between ANT and pulvinar (Supplementary Fig. 1).

Unlike in the thalamus, we noted a biphasic pattern in the group-level time course of SOZ aperiodic slope, with initial increase followed by a decrease below baseline levels at ∼60% of the ictal duration. SOZ slope has been reported to be decreased during seizures with respect to interictal baseline.^57^ Whereas SOZ slope was lower during the latter 40% (-0.72±0.47 *z*) vs. the first 60% (0.59±0.58 *z*) of the ictal duration (*z*=3.05, *P*=0.002, paired-sample Wilcoxon signed-rank test), SOZ slope in each of these epochs was no different from baseline (0-60%, *P*=0.5, *z=0.62*; 60-100%; *P*=0.1, *z*=1.55), although there was a trend towards aperiodic slope being decreased below baseline in the latter 40%.

### Local rhythms and beta-band inter-regional interactions link SOZ, near-SOZ, and thalamus

Seizures were associated with increased narrowband rhythmic activity in SOZ, near-SOZ, and thalamus. Ictal rhythms were characterized by identifying time-frequency (TF) clusters (*P*<0.05, permutation test, cluster-corrected) of spectral power (single subject example in Fig. 3A, group-level plots in Supplementary Fig. 2A). Consistent with the literature,^30,31,33,36^ we observed considerable diversity in their center frequencies: 10% of all clusters fell within the canonical delta (1-4 Hz) band; 16% in theta (4-8 Hz); 26% in alpha (8-13 Hz); 40% in beta (13-30 Hz); and 8% in gamma (30-150 Hz). We performed further analysis of ictal rhythms specifically within the alpha-beta frequency range, which comprised 66% of all TF clusters. Seizures were associated with increased alpha-beta power in all regions other than far-SOZ (*z*>3.51, *P*<10^-33^; Fig. 2E). Baseline-normalized alpha-beta power exceeded 2 z early in seizures, at latencies of: 14±3% in SOZ; 23±4% in near-SOZ; and 26±7% in thalamus (Fig. 2F).

**Figure 3:**
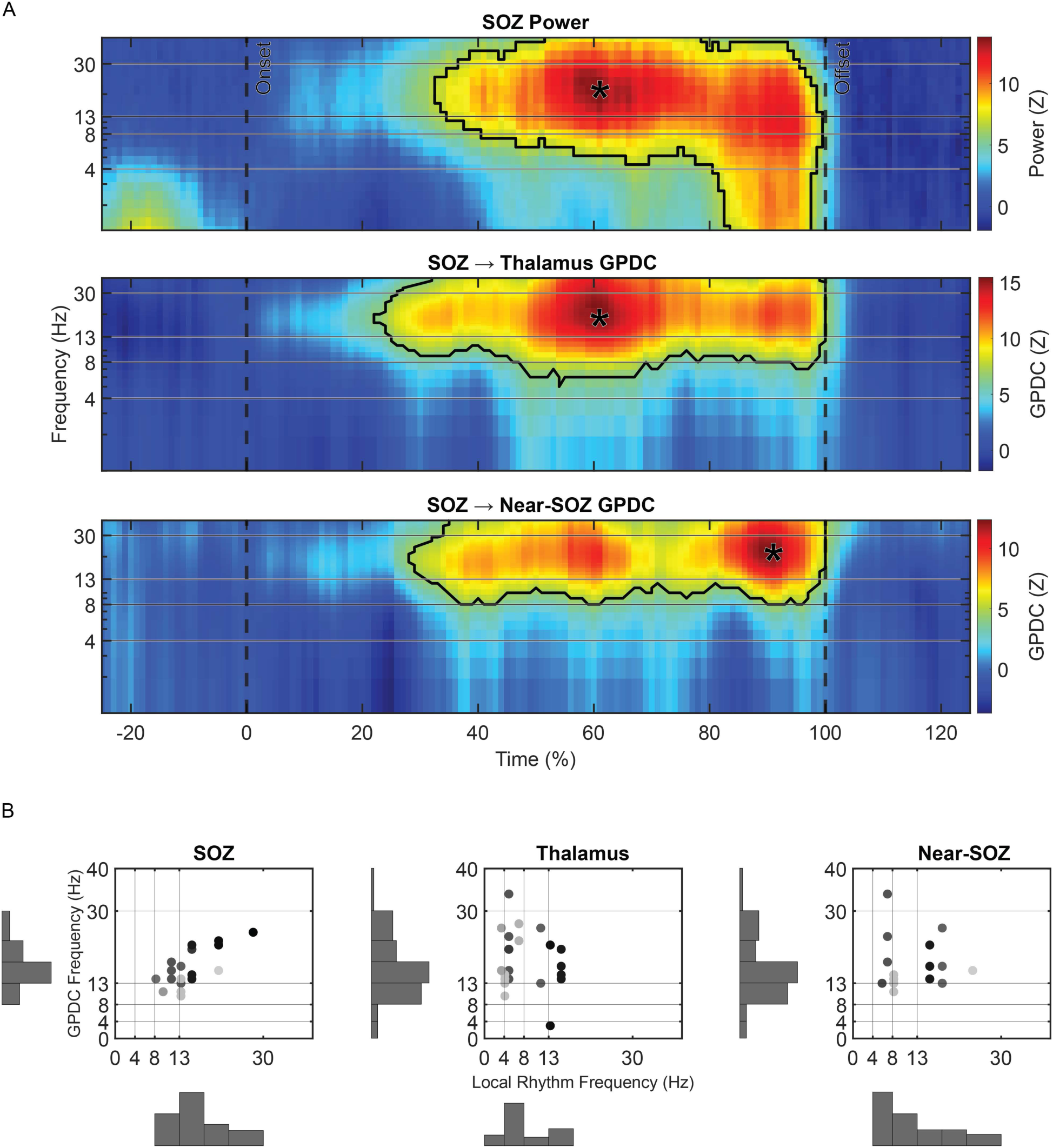
Inter-regional interactions throughout the thalamocortical ictal network involve beta band. **(A)** Example single-subject TF plots of spectral power (top panel) and GPDC strength (bottom two). Black outlines mark significant TF clusters (P<0.05, permutation test, cluster-corrected); asterisks mark their center frequencies. See Supplementary Fig. 2 for group-level results. **(B)** Center frequencies of local rhythms vs. inter-regional interactions. For each region, GPDC frequencies are shown for all other region pairs×directionality. Shadings mark individual subjects.

We next assessed inter-regional interactions using GPDC to obtain a frequency and direction-resolved measure of predictive (Granger) causality. Critically, GPDC is a multivariate metric that quantifies direct influence between region pairs while conditioning on the other regions included in the model. Seizures were associated with increased bidirectional interactions between SOZ, near-SOZ, and thalamus over interictal baseline levels (single-subject example in Fig. 3A, group-level results in Supplementary Fig. 2B). In contrast to local rhythms, inter-regional interactions involved the beta band in the majority of cases (90% of all significant TF clusters of GPDC strength). Notably, even in the few cases where the dominant local rhythms were outside of the beta band, significant GPDC clusters were centered in beta (Fig. 3B).

We further analyzed inter-regional interactions specifically within the beta band, confirming that seizures were associated with increased bidirectional GPDC between SOZ, near-SOZ, and thalamus (*z*>2.69, *P*<0.004; Fig. 4A). Note that interactions involving the far-SOZ were limited to outflow from the SOZ.

**Figure 4:**
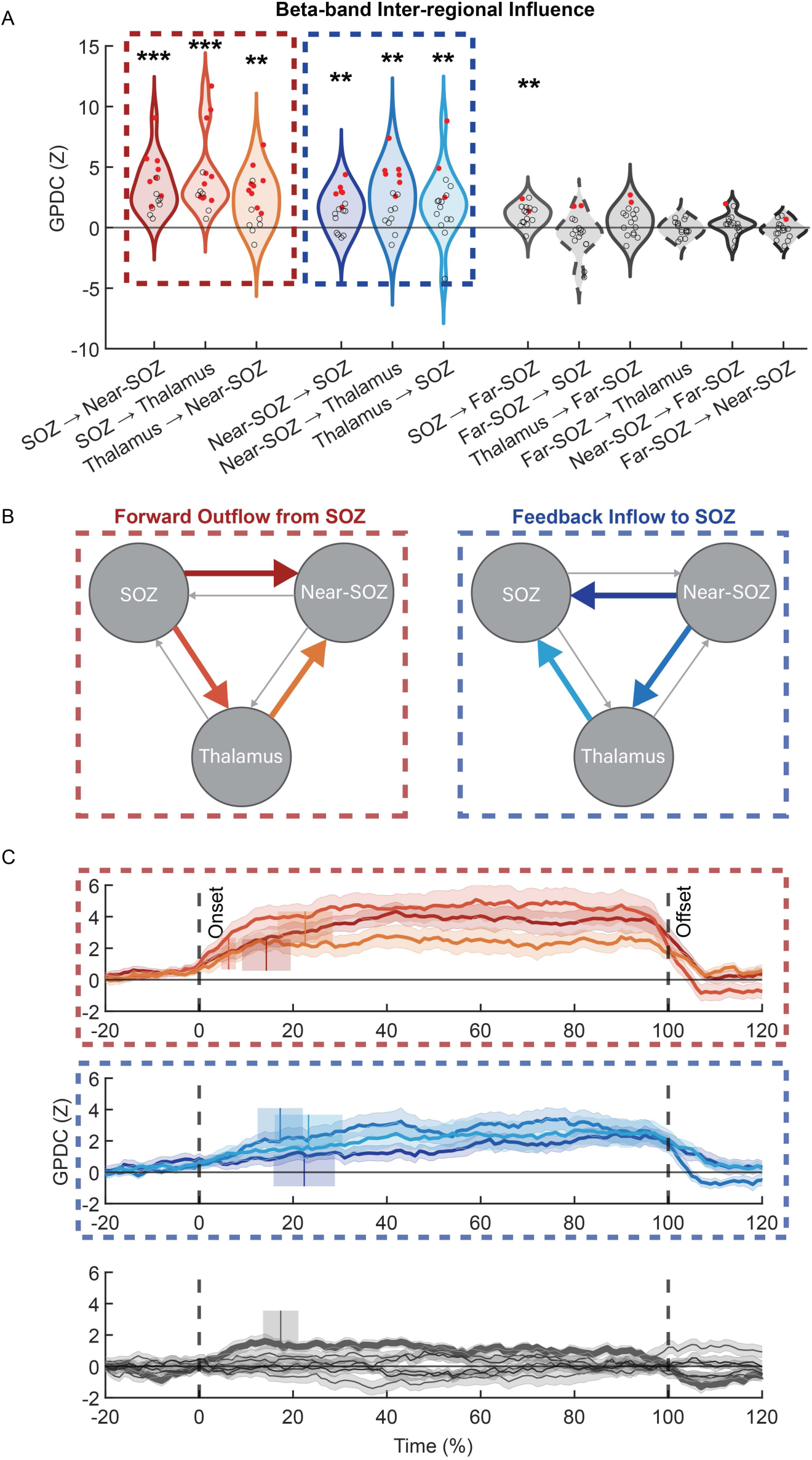
Bidirectional pathways of inter-regional interactions during seizures. GPDC interactions were grouped as follows: (i) forward outflow from SOZ (red outline), (ii) feedback inflow to SOZ (blue), and (iii) interactions involving the far-SOZ (shades of grey). **(A, B)** Time-averaged ictal beta-band GPDC strength (z-scored w.r.t. interictal baseline). Significance on subject-level tests marked in solid red (P<0.05, permutation test, cluster-corrected). Group-level significance assessed with one-sample Wilcoxon signed-rank test, FDR-corrected across region pairs×directionality. Both forward outflow from SOZ and feedback inflow to SOZ were expressed via a direct cortico-cortical pathway as well as an indirect pathway involving the thalamus (visually depicted in **B**). **(C)** Subject-averaged temporal dynamics of beta-band GPDC strength. In the bottom panel, time series for SOZ→far-SOZ marked with a thicker line. Vertical bars show average onset latency (>2 z w.r.t. baseline); shadings show s.e.m. \**P*<0.05, \*\**P*<0.01, \*\*\**P*<10^-3^. See Supplementary Fig. 1B for results stratified across distinct thalamic nuclei.

### Bidirectional pathways of ictal influence between SOZ and near-SOZ

The temporal dynamics of directed influence are shown in Fig. 4C, split into three groups as follows (see Fig. 4B for schematic): (i) forward outflow from SOZ (outlined in red in Fig 4), including a direct pathway (SOZ→near-SOZ) and an indirect pathway involving the thalamus (SOZ→thalamus, thalamus→near-SOZ); (ii) feedback inflow to SOZ (blue outline), again including direct (near-SOZ→SOZ) and indirect (near-SOZ→thalamus, thalamus→SOZ) pathways; (iii) interactions involving far-SOZ.

We first present findings on the temporal dynamics of forward outflow from the SOZ. Direct outflow (SOZ→near-SOZ) was increased (>2*z*) during early seizure at 14±5%. Indirect outflow was increased at 6±2% and 23%±6% for SOZ→thalamus and thalamus→near-SOZ, respectively. Direct comparisons showed that the onset latency of thalamic→near-SOZ followed that of SOZ→thalamus by 15%±6% (*z*=2.31, *P*=0.02). Interestingly, the influence of the SOZ on thalamus began earlier than its influence on near-SOZ (difference 9±5%; *z*=2.40, *P*=0.02). Moreover, the latencies of influence on near-SOZ through direct (SOZ→near-SOZ) vs. indirect pathways (thalamus→near-SOZ) were statistically indistinguishable (*z*=1.04, *P*=0.3). Together these findings highlight the prominent involvement of the thalamus in the early emergence of the ictal thalamocortical network. In terms of feedback to SOZ, direct inflow from near-SOZ was increased at 22±7%, whereas indirect inflow was increased at 17±5% and 23±7% for near-SOZ→thalamus and thalamus→SOZ, respectively. There was no difference between these latencies (*z*<1.02, *P*>0.3). We also compared latencies of forward vs. feedback interactions, and found no difference for either the direct or indirect pathways (*z*<1.34, *P*>0.2). In sum, our data show that not only ictal drive from SOZ, but also feedback influence on SOZ are constituents of *early* seizure dynamics.

Previous studies have reported that interactions between SOZ and thalamus ramp up leading up to seizure offset, specifically in the final 10-20 s.^25–28^. These findings have been interpreted as suggesting a role of the thalamus in seizure termination. Although not significant, our data showed consistent trends of increased GPDC during the early vs. late seizure epochs for both SOZ→thalamus (1.0±0.35 *z* vs. 1.5±0.45 *z*) and thalamus→SOZ interactions (0.28±0.25 *z* vs. 0.71±0.39 *z*; *z*<1.70, *P*>0.09). However, we observed that nearly all interactions between SOZ, near-SOZ, and thalamus ramped up throughout the duration of seizures (Supplementary Fig. 3). Previous studies have also reported that the dominant direction of interactions flips from SOZ→thalamus after onset to thalamus→SOZ before offset.^26,27^ In contrast, we found no difference in GPDC strengths between these two directions in either the early (SOZ→thalamus vs. thalamus→SOZ, 1.03±0.35 *z* vs. 0.28±0.25 *z*) or late epochs (1.47±0.45 *z* vs. 0.71±0.39 *z*; *z*<1.61, *P*>0.1). Rather, outflow from SOZ to thalamus trended towards being the dominant direction in both epochs.

### Thalamic aperiodic slope reflects cortical network interactions linked to seizure propagation

Having characterized ictal dynamics at both the local and network levels, we next sought to bridge between spatial scales. Motivated by recent models of the thalamocortical network by Jaramillo *et al*. showing that modulating the gain of thalamic neurons can bias the flow of neural activity between cortical regions,^42^ we tested the relationship between thalamic slope and the bidirectional interactions between SOZ and near-SOZ (Fig. 5A). Our analysis exploited the fact that while ictal dynamics showed consistent group-level patterns, the magnitude of ictal changes fluctuated across individual seizures (e.g., thalamic slope was decreased during seizures, but the size of this decrease varied across individual seizures). Specifically, we used a mixed-effects model with both within- and across-subject terms, and evaluated the regression coefficient of the within-subject term, which captures across-seizure associations between neural metrics that are consistently observed across subjects. On univariate analysis, we found that thalamic slope was not correlated with the magnitude of direct forward outflow (SOZ→near-SOZ; χ^2^=0.68, *P*=0.4).

**Figure 5:**
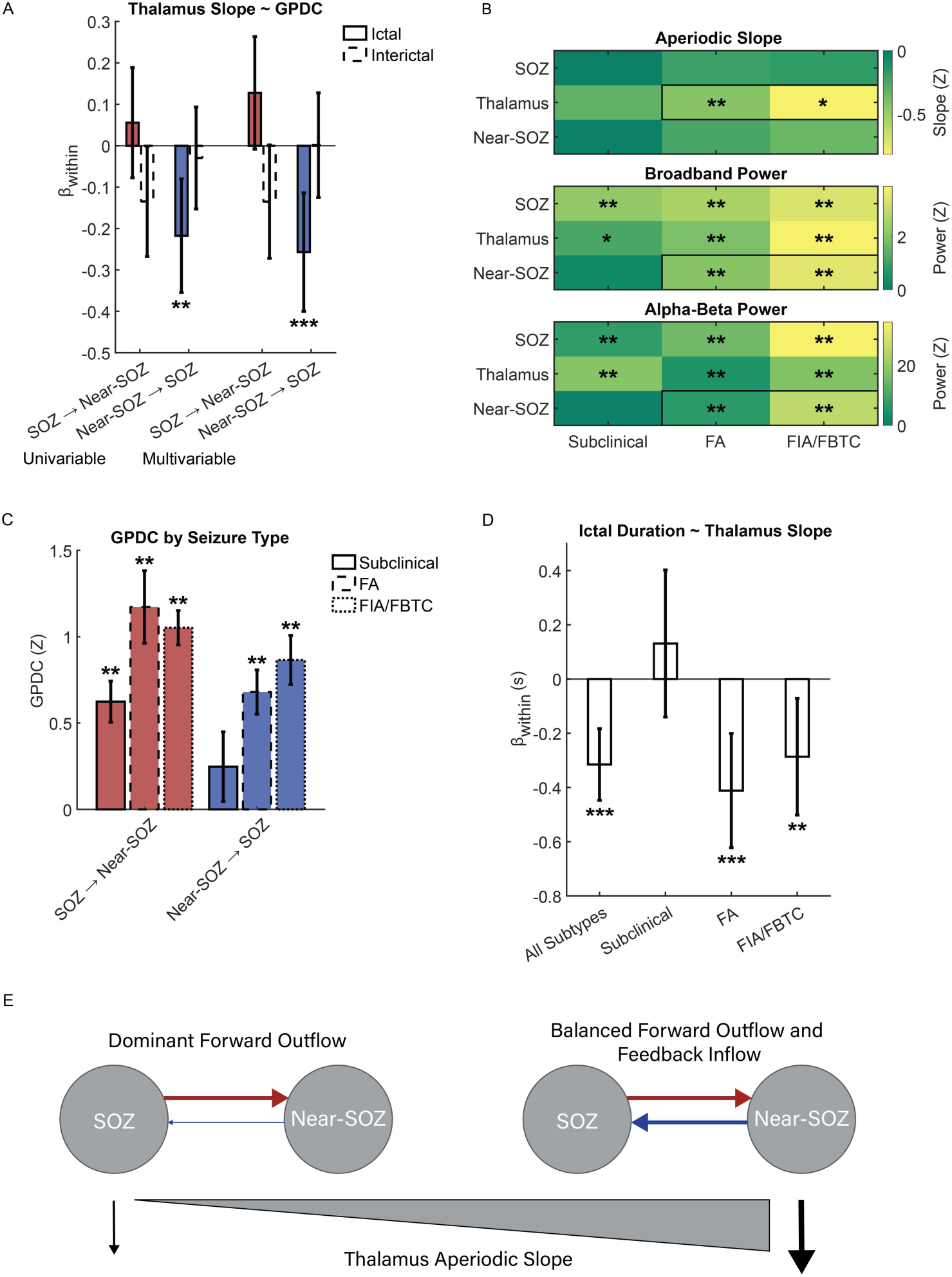
Ictal steepening of thalamic aperiodic slope corresponds to stronger feedback from near-SOZ to SOZ. **(A)** Seizure-to-seizure variability in the effect size of thalamic slope steepening is associated with the magnitude of feedback inflow (near-SOZ→SOZ; red bars), but not of forward outflow (SOZ→near-SOZ; blue). Bars show regression coefficient of within-subject term (within) from three models (solid outline): univariable models with (i) SOZ→near-SOZ terms or (ii) near-SOZ→SOZ terms; (iii) multivariable model with terms for both directions. Models constructed with interictal activity (dashed outline) are not significant. Error bars show 95% confidence intervals (C.I.). Significance assessed using Wald’s t-test, FDR-corrected across all tests, separately for ictal and interictal data. **(B)** Local ictal changes show distinct patterns across seizure types. Clinical seizures were distinguished by increased broadband and alpha-beta activity in near-SOZ (indicating seizure propagation), as well as decreased aperiodic slope in thalamus (outlined in black). In contrast, all seizure types showed increased broadband and alpha-beta activity in SOZ and thalamus. Colors represent subject and time-averaged values. Significance assessed with one-sample Wilcoxon signed-rank test, FDR-corrected across seizure types. **(C)** A similar pattern is observed at the network level. Forward outflow was increased in all seizure types, whereas feedback inflow was increased only in clinical seizures. Bars show subject-averaged GPDC strength; error bars show s.e.m. across subjects. Significance assessed with one-sample Wilcoxon signed-rank test, FDR-corrected across seizure types. **(D)** A more pronounced ictal steepening of thalamic slope predicted longer seizure durations. This effect was specific to the clinical seizure types. Significance assessed using Wald’s t-test, FDR-corrected across seizure types. **(E)** Diagram of cortical network states depending on the degree of thalamic slope steepening. FA: focal aware, FIA: focal impaired awareness; FBTC: focal to bilateral tonic clonic; \**P*<0.05, \*\**P*<0.01, \*\*\**P*<10^-3^.

In contrast, thalamic slope was negatively correlated with feedback inflow (near-SOZ→SOZ; χ^2^=9.52, *P*=0.002; β_*within*_=-0.22, *P*=0.002). This effect held in a model including GPDC terms for both directions, confirming that a larger ictal drop in thalamic slope is associated with greater feedback inflow from near-SOZ to SOZ (χ^2^*=12.89*, *P*=0.002; near-SOZ→SOZ, β_*within*_=-0.26, *P*=0.005; SOZ→near-SOZ, β_*within*_=0.13, *P*=0.07). On control analyses, identical models constructed with interictal activity were not significant (χ^2^<3.98, *P*>0.05).

We performed a similar analysis to examine how ictal changes in thalamic slope vary with *local* rhythmic activity, and found that fluctuations in thalamic slope were not associated with thalamic alpha-beta power across individual seizures (χ^2^=0.02, *P*=0.9). This stood in contrast to cortical aperiodic slope, which was negatively associated with local alpha-beta power across seizures in both SOZ and near-SOZ (χ^2^>9.14, *P*<0.002; β_*within*_<-5.2, *P*<0.003), recapitulating recent findings with intracranial recordings of physiologic dynamics.^58^

In short, our data revealed seizure-to-seizure covariation between thalamic slope and near-SOZ→SOZ feedback. Note that both of these dynamics showed significant deviations from interictal levels beginning in early seizure (22-23% after onset). To test their relationship to seizure propagation, we stratified further analysis across seizure types to capitalize on the fact that subclinical (electrographic) seizures typically do not propagate beyond the SOZ.^59^ Our data showed congruent findings of limited spread of ictal activity beyond the SOZ in subclinical seizures (Fig. 5B): both broadband and alpha-beta power were increased in near-SOZ in both clinical seizure subtypes (*z*>2.93, *P*<0.004), but not in subclinical seizures (*z*<0.42, *P*>0.5). Critically, thalamic slope demonstrated a similar pattern, whereby a significant drop was only observed in clinical seizures (*z*>2.37, *P*<0.02). Note that in contrast, thalamic broadband activity and low-frequency activity were increased in all seizure types (*z*>2.31, *P*<0.02). At the network level (Fig. 5C), whereas forward outflow from SOZ to near-SOZ was increased in all seizure types (*z*>2.67, *P*<0.008), significant increases in near-SOZ→SOZ feedback was again specific to clinical seizures (*z*>2.93, *P*<0.004).

In sum, as visually summarized in Fig. 5E, seizures with minimal change in thalamic slope coincided with cortical network states dominated by forward outflow from the SOZ. This pattern was characteristic of subclinical seizures, which had limited spread of ictal activity beyond the SOZ. Towards the other end of the spectrum, seizures with marked steepening of thalamic slope coincided with a more homogenous cortical network state, whereby the magnitudes of forward and feedback interactions between SOZ and near-SOZ were more balanced. This pattern distinguished clinical seizures, which showed propagation of seizure activity in near-SOZ. Hence, our data suggest that more prominent ictal steepening of thalamic aperiodic slope reflects network patterns that are linked to the propagation of seizures through the cortex. In a complementary analysis, we assessed for associations between thalamic slope and seizure duration using a similar modeling approach. We found converging evidence that the magnitude of thalamic slope steepening was negatively associated with seizure duration (χ^2^=21.19, *P*<10^-6^; β_*within*_=-0.32, *P*<10^-6^) – i.e., seizures with larger ictal steepening of thalamic slope were longer in duration (Fig. 5D). This held true in clinical seizures (χ^2^>6.68, *P*<0.01; β_*within*_<-0.28, *P*<0.01), but not in subclinical seizures (χ^2^=0.92, *P*>0.3).

## Discussion

### Ictal steepening of aperiodic slope is shared across thalamic nuclei

Seizures were associated with decreased (steeper) aperiodic slope in ANT and pulvinar, consistent with our prior work based on DBS recordings from ANT and CM nucleus.^36^ Collectively, these data show that ictal steepening of aperiodic slope is a thalamic signature shared across the nuclei that are most commonly targeted for neuromodulation. Our findings bridge to a broader literature showing that these very nuclei are involved in the coordination of brain-wide network dynamics,^40,41,60^ consistent with their widespread bidirectional anatomic connectivity with multiple cortical and limbic structures. Neuroimaging work has specifically implicated measures of neural variability^45^ (aperiodic slope is one such metric) in the thalamus as markers of brain-wide functional interactions across multiple timescales.^46,47^ Notably, prior intracranial studies of the thalamus in epilepsy have emphasized oscillatory activity, such that aperiodic activity remains an understudied feature of ictal thalamic activity. Here, we showed that thalamic aperiodic slope not only has dissociable dynamics from broadband and low-frequency activity, but also uniquely distinguished clinical from subclinical seizures.

### Ictal changes in cortical aperiodic slope

Previous intracranial studies (SEEG,^57^ device recordings^61,62^) have reported decreased aperiodic slope in the SOZ during seizures. Although our more granular analysis of the timecourse of cortical aperiodic slope showed its decline as seizures evolved, we did not find consistent ictal changes in cortical slope relative to baseline. We note the following methodologic differences: a narrower frequency range was used to compute aperiodic slope, following foundational methodologic studies;^54,63,64^ SEEG data allowed clear identification of seizure segments, and were not confounded by stimulation-induced activity.

Cortical slope was found to be flatter in generalized epilepsy vs. healthy controls.^65^ These data appeared to align with previous reports showing steeper slope during sleep and under anesthesia,^37–39^ which had led to the interpretation that a steeper slope corresponds to an E-I balance tipped towards inhibition. However, we note that the bulk of studies in epilepsy have reported seemingly contradictory findings: aperiodic slope is flatter in non-SOZ vs. SOZ;^57^ flattening of cortical slope tracks favorable therapeutic responses to drugs^66^ and neuromodulation.^61,67–69^

### Local and network rhythms throughout the thalamocortical ictal network

Ictal rhythms in the thalamus, as in the cortex, are heterogeneous in their frequency content, spanning both lower and higher frequency bands.^30,31,33,36^ In contrast, we showed that inter-regional interactions across the thalamocortical network primarily involved the beta band. Interestingly, cortical beta power has been shown to track the cortical spread of seizure activity in animal models.^70^ The majority of prior studies on thalamocortical functional connectivity have not assessed the frequency specificity of regional interactions.^25,27–29^ Panchavati *et al*. analyzed early (first 20 s) seizure dynamics, and showed that bidirectional interactions between SOZ and thalamus (ANT, CM) spanned the canonical frequency bands, from delta to gamma.^32^ Damiani *et al*. similarly observed interactions involving all of the lower bands (theta to beta), although beta-band interactions demonstrated the highest regional specificity across cortical regions stratified based on connectivity with the thalamus.^26^

### Thalamic aperiodic slope reflects cortical network dynamics linked to seizure propagation

Whereas cortical aperiodic slope was associated with local low-frequency rhythms, thalamic aperiodic slope reflected network-level patterns in beta-band inter-regional interactions. We found that steeper thalamic aperiodic slope corresponded to cortical network states with more balanced interactions between the SOZ and near-SOZ. This pattern was characteristic of clinical seizures, which also showed propagated seizure activity in the near-SOZ. Together these results suggest that a more homogenous network topology in the cortex may be conducive to seizure spread, and that the thalamus might be involved in this process. We emphasize that increased Granger causality *per se* (as well as synchrony) do not indicate whether the net effects on the target region are excitatory/activating vs. inhibitory/suppressive. In this section, we discuss these interpretations in light of previous literature.

From a broader perspective, there is growing evidence to indicate a critical role of the thalamus in shaping cortex-wide neural dynamics.^40,41^ Using biologically-plausible models of thalamocortical networks, Jaramillo *et al*. showed that the flow of cortical activity can be dynamically controlled by the local activity of the thalamus.^42^ In their model, the key variable was the excitability of the thalamus, which was operationalized by the gain of thalamic neurons. Our findings of a steeper thalamic aperiodic slope during seizures may indicate an E-I balance tipped towards the inhibition,^37–39^ which is thought to correspond to decreased neuronal gain. Our results confirm two model predictions in terms of the expected effects of steeper thalamic slope/decreased thalamic gain on cortical network dynamics: increased symmetry of inter-regional functional connectivity in the cortex (i.e., a more homogenous functional architecture), and increased low-frequency oscillations both within and between cortical regions.

Network-level studies of cortical ictal dynamics have suggested that non-SOZ regions play an important role in the cortical propagation of seizures. Khambhati *et al*. showed that seizures that do not generalize (FA, FIA) were distinguished from FBTC seizures by a preponderance of non-SOZ nodes that influence the synchronizability of the cortical network.^20^ Interestingly, both synchronizing and desynchronizing nodes in the non-SOZ were more common in seizures that remained focal. This suggests that increased network heterogeneity may limit seizure spread, which aligns with our data.

In contrast to our results, Jiang *et al*. showed that non-SOZ→SOZ influence was greater in seizures that did not generalize, whereas there was no difference in the magnitude of SOZ→non-SOZ influence.^18^ We note the following methodological concerns that may account for the discrepant findings. First, the authors included all non-SOZ channels in their functional connectivity model, and took the average across all pairs of SOZ and non-SOZ channels to quantify the aggregate influence of the non-SOZ on SOZ. This assumes that the non-SOZ is functionally uniform in terms of its participation in seizure dynamics. However, our data as well as Khambhati *et al*.^20^ demonstrate the heterogeneous contributions of the non-SOZ. Second, Jiang *et al*. utilized the directed transfer function (DTF), which (unlike PDC/GPDC) does not distinguish direct pathways of influence from cascade pathways (influence relayed through other modeled nodes/channels). Therefore, it is possible that the reported influence of non-SOZ on SOZ may have simply been relayed by other nodes within the SOZ.

In a more recent study, Makhoul *et al*. did account for the heterogeneity of the non-SOZ by stratifying into an “early propagation zone” (defined by seizure spread within 10 s of onset) and a “non-involved zone.”^21^ Notably, in spite of the label, the “non-involved zone” showed propagated seizure activity within 10-20 s of onset (see Fig. 4B). The authors reported two distinct phases of cortical seizure dynamics. In the first phase, inflow was increased from the “non-involved zone” to the SOZ and “early propagation zone.” In the second phase, this inflow dropped in FIA and FBTC seizures, but not in FA seizures, which was interpreted as loss of the suppressive effects of the “non-involved zone,” permitting seizure spread. We note the following methodological differences from our study. First, Makhoul *et al*. utilized PDC, which unlike GPDC is more susceptible to spurious estimates driven by cross-channel differences in prediction error (additional details in **Materials and methods**). Second, how the loss of inflow from the “non-involved zone” in FIA/FBTC seizures ultimately impacted the cortical extent of seizure spread was not explicitly shown. In contrast, we demonstrated the relative lack of propagated ictal activity in the near-SOZ in subclinical vs. clinical seizures, allowing us to more clearly interpret the implications of increased near-SOZ→SOZ feedback on seizure propagation.

In sum, the literature to date suggests that interplay between SOZ and non-SOZ may be critical to seizure propagation, although the specific dynamics remain to be further elucidated. Our findings add to this literature, indicating that the thalamus may be involved in biasing inter-regional interactions in the cortex towards network states that may facilitate seizure spread. These results dovetail nicely with a previous intracranial study showing that seizures with early onset of fast ictal activity in the thalamus (as quantified by the epileptogenicity index) also had spatially-larger extents of elevated epileptogenicity index in the cortex.^34^

### A role of the thalamus in seizure termination?

In a series of studies by Bartolomei and colleagues, and more recently by Damiani *et al*., thalamocortical interactions ramped up in magnitude leading up to offset.^25–28^ The dominant direction of thalamocortical interactions was reported to reverse directions, from cortex→thalamus to thalamus→cortex.^26,27^ Greater thalamic outflow towards the end of seizures was furthermore associated with seizures exhibiting synchronous cortical offset,^28^ as well as seizures with terminal electrographic patterns indicative of strong thalamocortical synchrony.^25^ Together these findings have been interpreted as suggesting a role of the thalamus in seizure termination. Although not significant, our data revealed consistent trends showing that interactions between SOZ and thalamus (albeit in both directions) were greater in late vs. early seizure. However, SOZ→thalamus influence trended towards being greater vs. the reverse direction in both epochs. We note several methodological strengths of our work: thalamocortical interactions were stratified with respect to cortices within or outside the SOZ; inter-regional interactions were quantified with a multivariate method.

### Limitations

To analyze the generalized dynamics of seizures requires some amount of averaging across seizures and subjects. Methodologic challenges stem from the inherent spatial and temporal variability of seizures, both within and across subjects. To control for variable seizure durations, we utilized a linear normalization to map time to percentages spanning each individual seizure duration (as in ref^31^). We acknowledge that this approach assumes linearity of temporal dynamics across seizures. An alternative is to analyze average dynamics in raw time, which has been done to study initiation and termination dynamics.^25–27,29,30,32^ However, this requires choosing arbitrary cutoff values (studies have focused on the first/last 10-20 s), and further assumes that initiation and termination dynamics are stereotyped across seizures, which is not necessarily true.^71,72^ Non-linear methods (e.g., dynamic time warping) may provide complementary insights, although they are not devoid of assumptions in constructing a distance function that is minimized to align signals. We analyzed inter-regional interactions using a multivariate approach, explicitly stratifying the non-SOZ into near- and far-SOZ. Nonetheless, we acknowledge that the heterogeneous contributions of the non-SOZ to seizure dynamics may require an even more spatially-granular analysis. We further acknowledge that our results on the latencies and directionality of seizure dynamics do not imply causation *per se*. Alternative approaches using effective connectivity (e.g., dynamic causal modeling) may offer converging evidence, although these methods do not directly demonstrate causality either. Finally, we note that correlation between standard visual assessments of seizure propagation to thalamic nuclei and quantitative metrics was outside the scope of this study.

### Conclusions

The thalamus was rapidly recruited in the emergence of the ictal network, bound together by bidirectional interactions between the SOZ, thalamus, and cortices surrounding the SOZ (near-SOZ). Seizures were associated with an early, sustained drop in aperiodic slope that was regionally specific to the thalamus. More pronounced thalamic slope steepening was associated with stronger near-SOZ→SOZ feedback across individual seizures. Comparisons of these dynamics between subclinical and clinical seizures was revealing, leveraging the fact that clinical seizures, but not subclinical seizures, showed propagated ictal activity in near-SOZ. We found that only clinical seizures showed significant steepening of thalamic slope, as well as near-SOZ→SOZ feedback that was increased to levels commensurate with SOZ→near-SOZ outflow. Together these findings suggest a putative role of the thalamus in the spatial propagation of seizures across the cortex, expanding on prior intracranial work emphasizing its possible involvement in seizure termination.^25–28^ More broadly, these results align with a growing literature implicating the thalamus in the coordination and regulation of cortico-cortical interactions.^40,41,60,73^ Our data further highlights thalamic aperiodic slope as a candidate physiologic index of cortical states linked to seizure spread, with potential relevance to physiology-guided neuromodulation.

## Data Availability

Data may be made available upon reasonable request.

## Acknowledgements

We are deeply grateful to the patients who gave consent for the use of their SEEG data for the advancement of knowledge and the benefit of future epilepsy patients. We also thank the clinical staff in the Epilepsy Monitoring Unit for their dedication to patient care, and members of Barrow Informatics and Neuromodulation Research for their assistance with data collection and technical support. Large language models were used for grammar review of the manuscript; minimal edits that did not change the content of the manuscript were incorporated.

---

**Supplementary Figure 1:**
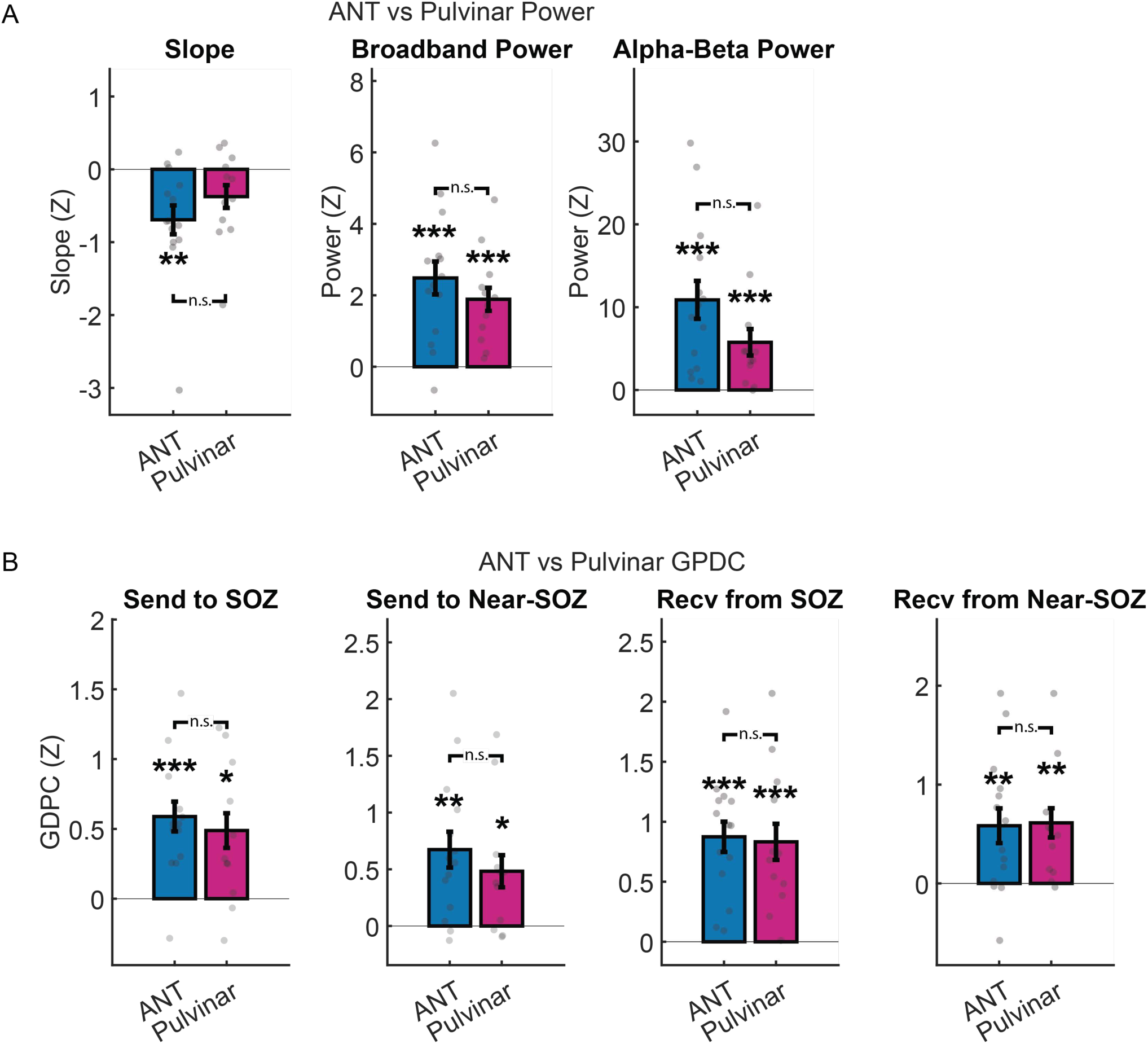
Ictal dynamics were consistent across ANT (209 seizures in 14 subjects) and pulvinar (146 seizures in 10 subjects), both in terms of their **(A)** local activity as well as in their **(B)** interactions with cortical ROIs. Group-level significance assessed with one-sample Wilcoxon signed-rank test, FDR-corrected across thalamic nuclei. Comparisons across thalamic nuclei assess with paired-sample Wilcoxon signed-rank test. n.s.: not significant; \**P*<0.05, \*\**P*<0.01, \*\*\**P*<10^-3^.

**Supplementary Figure 2:**
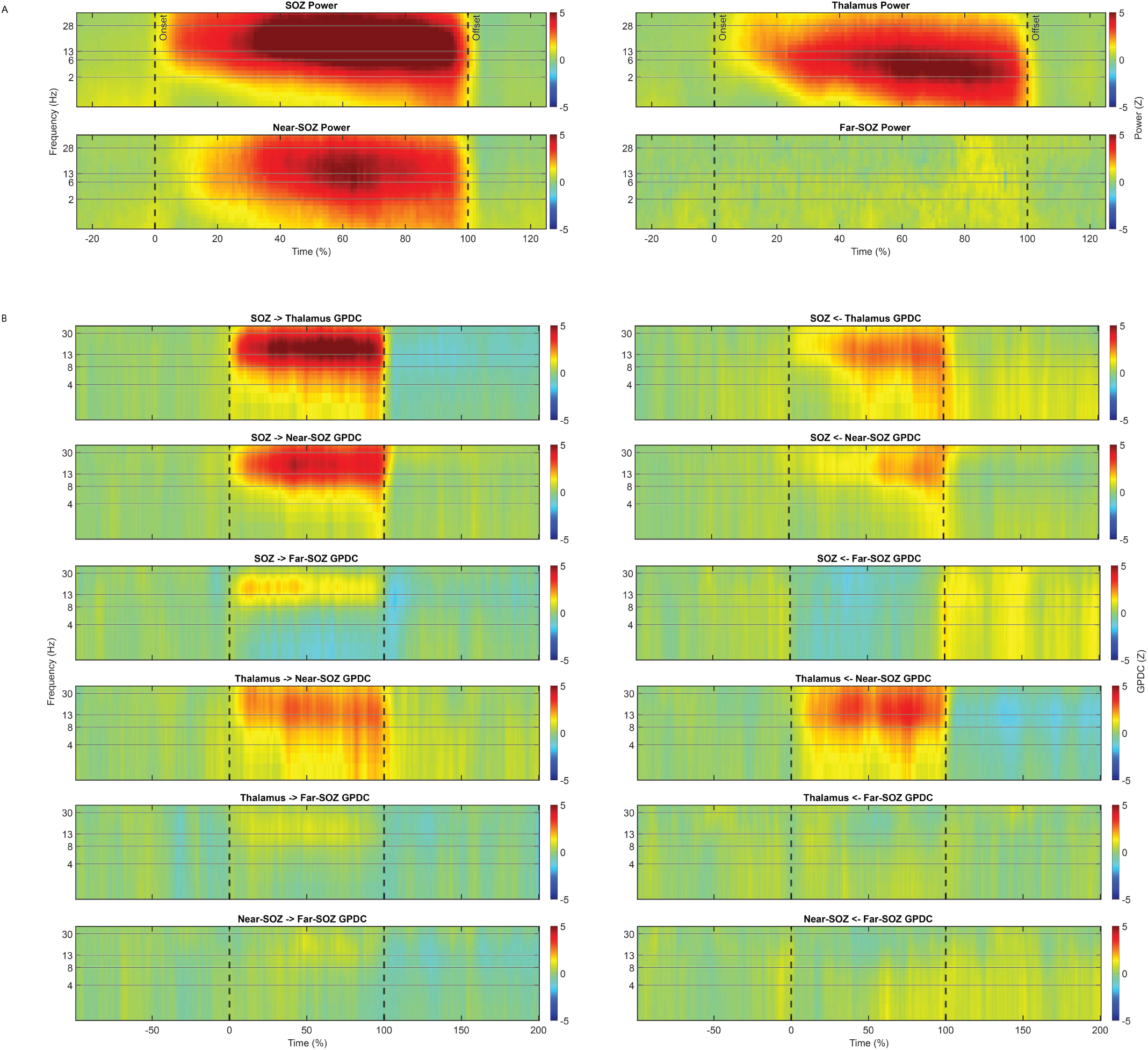
Group-level time-frequency plots showing subject-averaged values of **(A)** spectral power and **(B)** GPDC strengths.

**Supplementary Figure 3:**
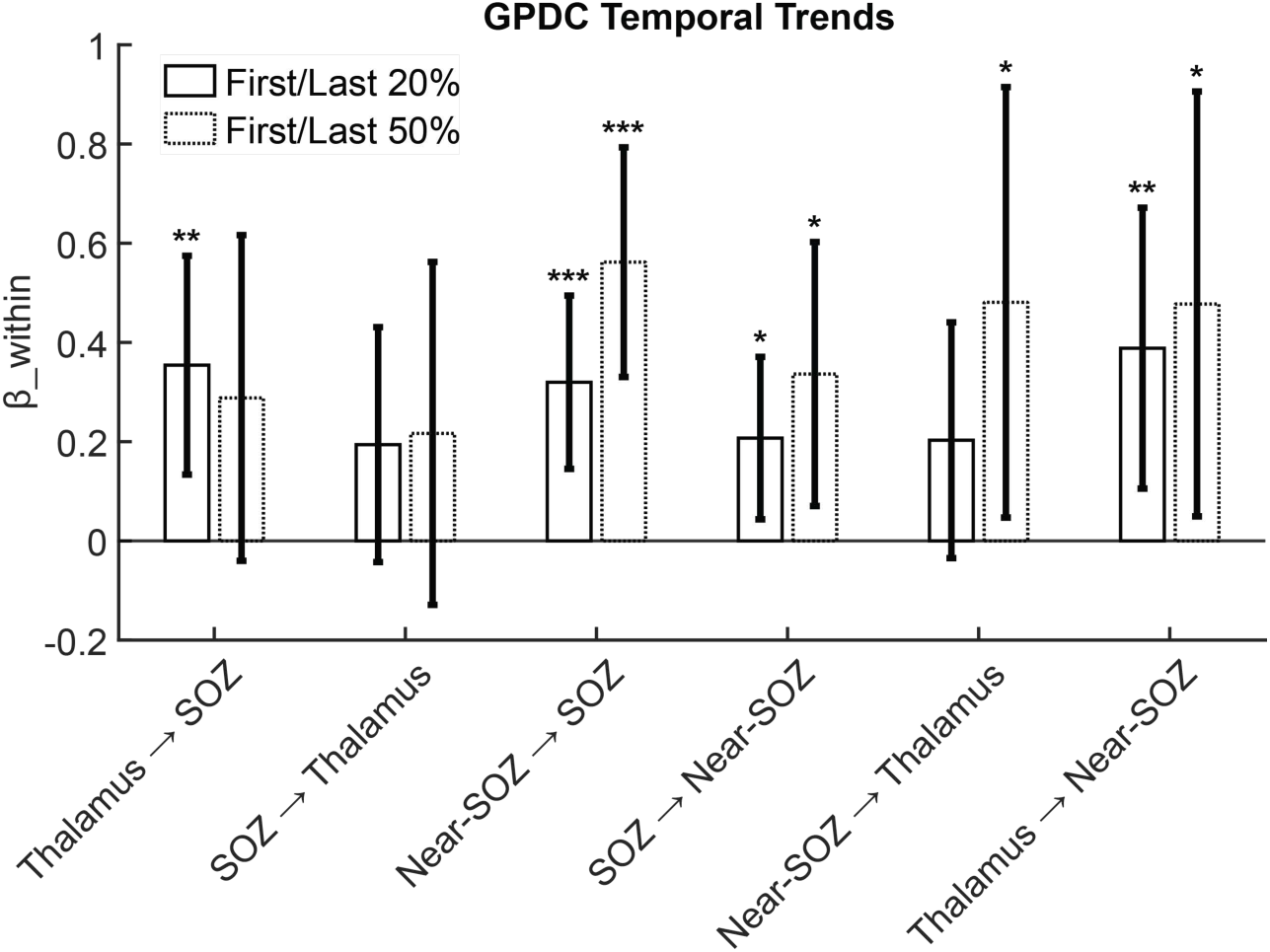
GPDC strengths ramped up during seizures for all region pairs and directions except SOZ→thalamus. Mixed-effects models were used to compare between the first and last 20% (solid outline) or the first and second 50% of the seizure duration (dotted), separately for each region pair×directionality. Bars show regression coefficients of models; error bars show 95% C.I. Significance assessed using Wald’s t-test, FDR-corrected across all models within each analysis epoch pair. n.s.: not significant; \**P*<0.05, \*\**P*<0.01, \*\*\**P*<10^-3^.

## References

1. Fisher R, Salanova V, Witt T, et al. Electrical stimulation of the anterior nucleus of thalamus for treatment of refractory epilepsy. Epilepsia. 2010;51(5):899–908. doi:10.1111/j.1528-1167.2010.02536.x

2. Salanova V, Sperling MR, Gross RE, et al. The SANTÉ study at 10 years of follow-up: Effectiveness, safety, and sudden unexpected death in epilepsy. Epilepsia. 2021;62(6):1306–1317. doi:10.1111/epi.16895

3. Peltola J, Colon AJ, Pimentel J, et al. Deep Brain Stimulation of the Anterior Nucleus of the Thalamus in Drug-Resistant Epilepsy in the MORE Multicenter Patient Registry. Neurology. 2023;100(18):e1852–e1865. doi:10.1212/WNL.0000000000206887

4. Dalic LJ, Warren AEL, Bulluss KJ, et al. DBS of Thalamic Centromedian Nucleus for Lennox-Gastaut Syndrome (ESTEL Trial). Ann Neurol. 2022;91(2):253–267. doi:10.1002/ana.26280

5. Pizzo F, Carron R, Laguitton V, Clement A, Giusiano B, Bartolomei F. Medial pulvinar stimulation for focal drug-resistant epilepsy: interim 12-month results of the PULSE study. Front Neurol. 2024;15:1480819. doi:10.3389/fneur.2024.1480819

6. Li MCH, Cook MJ. Deep brain stimulation for drug-resistant epilepsy. Epilepsia. 2018;59(2):273–290. doi:10.1111/epi.13964

7. Yang AI, Isbaine F, Alwaki A, Gross RE. Multitarget deep brain stimulation for epilepsy. Journal of Neurosurgery. 2023;1(aop):1–8. doi:10.3171/2023.5.JNS23982

8. McGinn R, Von Stein EL, Datta A, et al. Ictal Involvement of the Pulvinar and the Anterior Nucleus of the Thalamus in Patients With Refractory Epilepsy. Neurology. 2024;103(11):e210039. doi:10.1212/WNL.0000000000210039

9. Wu TQ, Kaboodvand N, McGinn RJ, et al. Multisite thalamic recordings to characterize seizure propagation in the human brain. Brain. 2023;146(7):2792–2802. doi:10.1093/brain/awad121

10. Gadot R, Korst G, Shofty B, Gavvala JR, Sheth SA. Thalamic stereoelectroencephalography in epilepsy surgery: a scoping literature review. J Neurosurg. Published online March 11, 2022:1–16. doi:10.3171/2022.1.JNS212613

11. Carron R, Pizzo F, Trébuchon A, Bartolomei F. Letter to the Editor. Thalamic sEEG and epilepsy. J Neurosurg. 2022;138(4):1172–1173. doi:10.3171/2022.9.JNS222169

12. Wang K, Zhang X, Song C, et al. Decreased Intrinsic Neural Timescales in Mesial Temporal Lobe Epilepsy. Front Hum Neurosci. 2021;15:772365. doi:10.3389/fnhum.2021.772365

13. Xie K, Royer J, Lariviere S, et al. Atypical intrinsic neural timescales in temporal lobe epilepsy. Epilepsia. 2023;64(4):998–1011. doi:10.1111/epi.17541

14. Royer J, Larivière S, Rodriguez-Cruces R, et al. Cortical microstructural gradients capture memory network reorganization in temporal lobe epilepsy. Brain. 2023;146(9):3923–3937. doi:10.1093/brain/awad125

15. Spencer SS. Neural networks in human epilepsy: evidence of and implications for treatment. Epilepsia. 2002;43(3):219–227. doi:10.1046/j.1528-1157.2002.26901.x

16. Piper RJ, Richardson RM, Worrell G, et al. Towards network-guided neuromodulation for epilepsy. Brain. 2022;145(10):3347–3362. doi:10.1093/brain/awac234

17. Laufs H. Functional imaging of seizures and epilepsy: evolution from zones to networks. Curr Opin Neurol. 2012;25(2):194–200. doi:10.1097/WCO.0b013e3283515db9

18. Jiang H, Cai Z, Worrell GA, He B. Multiple Oscillatory Push–Pull Antagonisms Constrain Seizure Propagation. Annals of Neurology. 2019;86(5):683–694. doi:10.1002/ana.25583

19. Karimi-Rouzbahani H, McGonigal A. Directionality of neural activity in and out of the seizure onset zone in focal epilepsy. Netw Neurosci. 2025;9(2):798–823. doi:10.1162/netn_a_00454

20. Khambhati AN, Davis KA, Lucas TH, Litt B, Bassett DS. Virtual Cortical Resection Reveals Push-Pull Network Control Preceding Seizure Evolution. Neuron. 2016;91(5):1170–1182. doi:10.1016/j.neuron.2016.07.039

21. Makhoul GS, Doss DJ, Johnson GW, et al. Collapse of interictal suppressive networks permits seizure spread. Brain. 2025;148(12):4275–4287. doi:10.1093/brain/awaf215

22. Liu Z, Han F, Yu Y, Wang Q. Role of hierarchical heterogeneity in shaping seizure onset dynamics: Insights from structurally-based whole-brain dynamical network models. Communications in Nonlinear Science and Numerical Simulation. 2024;130:107721. doi:10.1016/j.cnsns.2023.107721

23. Meeren HKM, Pijn JPM, Van Luijtelaar ELJM, Coenen AML, Lopes da Silva FH. Cortical focus drives widespread corticothalamic networks during spontaneous absence seizures in rats. J Neurosci. 2002;22(4):1480–1495. doi:10.1523/JNEUROSCI.22-04-01480.2002

24. McCormick DA, Contreras D. On the cellular and network bases of epileptic seizures. Annu Rev Physiol. 2001;63:815–846. doi:10.1146/annurev.physiol.63.1.815

25. Evangelista E, Bénar C, Bonini F, et al. Does the Thalamo-Cortical Synchrony Play a Role in Seizure Termination? Front Neurol. 2015;6:192. doi:10.3389/fneur.2015.00192

26. Damiani A, Nouduri S, Ho JC, et al. Thalamocortical hodology to personalize electrical stimulation for focal epilepsy. Nat Commun. 2025;16(1):9209. doi:10.1038/s41467-025-64922-w

27. Guye M, Régis J, Tamura M, et al. The role of corticothalamic coupling in human temporal lobe epilepsy. Brain. 2006;129(Pt 7):1917–1928. doi:10.1093/brain/awl151

28. Soulier H, Pizzo F, Jegou A, et al. The anterior and pulvinar thalamic nuclei interactions in mesial temporal lobe seizure networks. Clin Neurophysiol. 2023;150:176–183. doi:10.1016/j.clinph.2023.03.016

29. Arthuis M, Valton L, Régis J, et al. Impaired consciousness during temporal lobe seizures is related to increased long-distance cortical-subcortical synchronization. Brain. 2009;132(Pt 8):2091–2101. doi:10.1093/brain/awp086

30. Pizarro D, Ilyas A, Chaitanya G, et al. Spectral organization of focal seizures within the thalamotemporal network. Ann Clin Transl Neurol. 2019;6(9):1836–1848. doi:10.1002/acn3.50880

31. Ilyas A, Toth E, Chaitanya G, Riley K, Pati S. Ictal high-frequency activity in limbic thalamic nuclei varies with electrographic seizure-onset patterns in temporal lobe epilepsy. Clin Neurophysiol. 2022;137:183–192. doi:10.1016/j.clinph.2022.01.134

32. Panchavati S, Daida A, Edmonds B, et al. Uncovering spatiotemporal dynamics of the corticothalamic network at ictal onset. Epilepsia. 2024;65(7):1989–2003. doi:10.1111/epi.17990

33. Singh J, Miller JA, Lucas T, et al. Anterior thalamic nucleus local field potentials during focal temporal lobe epileptic seizures. Front Neurol. 2024;15:1419835. doi:10.3389/fneur.2024.1419835

34. Pizzo F, Roehri N, Giusiano B, et al. The Ictal Signature of Thalamus and Basal Ganglia in Focal Epilepsy: A SEEG Study. Neurology. 2021;96(2):e280–e293. doi:10.1212/WNL.0000000000011003

35. Ilyas A, Alamoudi OA, Riley KO, Pati S. Pro-Ictal State in Human Temporal Lobe Epilepsy. NEJM Evidence. 2023;2(3):EVIDoa2200187. doi:10.1056/EVIDoa2200187

36. Yang AI, Raghu ALB, Isbaine F, Alwaki A, Gross RE. Sensing with deep brain stimulation device in epilepsy: aperiodic changes in thalamic local field potential during seizures. Epilepsia. Published online August 22, 2023. doi:10.1111/epi.17758

37. Waschke L, Donoghue T, Fiedler L, et al. Modality-specific tracking of attention and sensory statistics in the human electrophysiological spectral exponent. Elife. 2021;10:e70068. doi:10.7554/eLife.70068

38. Lendner JD, Helfrich RF, Mander BA, et al. An electrophysiological marker of arousal level in humans. Haegens S, Colgin LL, Piantoni G, eds. eLife. 2020;9:e55092. doi:10.7554/eLife.55092

39. Gao R, Peterson EJ, Voytek B. Inferring synaptic excitation/inhibition balance from field potentials. Neuroimage. 2017;158:70–78. doi:10.1016/j.neuroimage.2017.06.078

40. Hwang K, Bertolero MA, Liu WB, D’Esposito M. The Human Thalamus Is an Integrative Hub for Functional Brain Networks. J Neurosci. 2017;37(23):5594–5607. doi:10.1523/JNEUROSCI.0067-17.2017

41. Halassa MM, Kastner S. Thalamic functions in distributed cognitive control. Nat Neurosci. 2017;20(12):1669–1679. doi:10.1038/s41593-017-0020-1

42. Jaramillo J, Mejias JF, Wang XJ. Engagement of Pulvino-cortical Feedforward and Feedback Pathways in Cognitive Computations. Neuron. 2019;101(2):321–336.e9. doi:10.1016/j.neuron.2018.11.023

43. Chance FS, Abbott LF, Reyes AD. Gain modulation from background synaptic input. Neuron. 2002;35(4):773–782. doi:10.1016/s0896-6273(02)00820-6

44. Bhatia A, Moza S, Bhalla US. Precise excitation-inhibition balance controls gain and timing in the hippocampus. eLife. 8:e43415. doi:10.7554/eLife.43415

45. Waschke L, Kloosterman NA, Obleser J, Garrett DD. Behavior needs neural variability. Neuron. 2021;109(5):751–766. doi:10.1016/j.neuron.2021.01.023

46. Garrett DD, Epp SM, Perry A, Lindenberger U. Local temporal variability reflects functional integration in the human brain. NeuroImage. 2018;183:776–787. doi:10.1016/j.neuroimage.2018.08.019

47. Garrett DD, Skowron A, Wiegert S, et al. Lost Dynamics and the Dynamics of Loss: Longitudinal Compression of Brain Signal Variability is Coupled with Declines in Functional Integration and Cognitive Performance. Cereb Cortex. 2021;31(11):5239–5252. doi:10.1093/cercor/bhab154

48. Manning JR, Jacobs J, Fried I, Kahana MJ. Broadband shifts in local field potential power spectra are correlated with single-neuron spiking in humans. J Neurosci. 2009;29(43):13613–13620. doi:10.1523/JNEUROSCI.2041-09.2009

49. Dale AM, Fischl B, Sereno MI. Cortical surface-based analysis. I. Segmentation and surface reconstruction. Neuroimage. 1999;9(2):179-194. doi:10.1006/nimg.1998.0395

50. Fischl B, Salat DH, Busa E, et al. Whole brain segmentation: automated labeling of neuroanatomical structures in the human brain. Neuron. 2002;33(3):341–355. doi:10.1016/s0896-6273(02)00569-x

51. Iglesias JE, Insausti R, Lerma-Usabiaga G, et al. A probabilistic atlas of the human thalamic nuclei combining ex vivo MRI and histology. Neuroimage. 2018;183:314–326. doi:10.1016/j.neuroimage.2018.08.012

52. Desikan RS, Ségonne F, Fischl B, et al. An automated labeling system for subdividing the human cerebral cortex on MRI scans into gyral based regions of interest. Neuroimage. 2006;31(3):968–980. doi:10.1016/j.neuroimage.2006.01.021

53. Mazziotta J, Toga A, Evans A, et al. A probabilistic atlas and reference system for the human brain: International Consortium for Brain Mapping (ICBM). Philos Trans R Soc Lond B Biol Sci. 2001;356(1412):1293–1322. doi:10.1098/rstb.2001.0915

54. Donoghue T, Haller M, Peterson EJ, et al. Parameterizing neural power spectra into periodic and aperiodic components. Nature Neuroscience. 2020;23(12):12. doi:10.1038/s41593-020-00744-x

55. Baccala LA, Sameshima K, Takahashi DY. Generalized Partial Directed Coherence. In: 2007 15th International Conference on Digital Signal Processing. 2007:163–166. doi:10.1109/ICDSP.2007.4288544

56. Maris E, Oostenveld R. Nonparametric statistical testing of EEG- and MEG-data. J Neurosci Methods. 2007;164(1):177–190. doi:10.1016/j.jneumeth.2007.03.024

57. Jiang H, Kokkinos V, Ye S, et al. Interictal SEEG Resting-State Connectivity Localizes the Seizure Onset Zone and Predicts Seizure Outcome. Adv Sci (Weinh). 2022;9(18):e2200887. doi:10.1002/advs.202200887

58. Preston M, Schaworonkow N, Voytek B. Time-Resolved Aperiodic and Oscillatory Dynamics during Human Visual Memory Encoding. J Neurosci. 2025;45(16):e2404242025. doi:10.1523/JNEUROSCI.2404-24.2025

59. Zangaladze A, Nei M, Liporace JD, Sperling MR. Characteristics and clinical significance of subclinical seizures. Epilepsia. 2008;49(12):2016–2021. doi:10.1111/j.1528-1167.2008.01672.x

60. Shine JM, Lewis LD, Garrett DD, Hwang K. The impact of the human thalamus on brain-wide information processing. Nat Rev Neurosci. 2023;24(7):416–430. doi:10.1038/s41583-023-00701-0

61. Satzer D, Kaye LC, Ojemann SG, Kramer DR, Thompson JA. Aperiodic activity as a biomarker of seizures and neuromodulation. Brain Stimul. 2025;18(3):738–744. doi:10.1016/j.brs.2025.03.022

62. Charlebois CM, Anderson DN, Smith EH, et al. Circadian changes in aperiodic activity are correlated with seizure reduction in patients with mesial temporal lobe epilepsy treated with responsive neurostimulation. Epilepsia. 2024;65(5):1360–1373. doi:10.1111/epi.17938

63. Janjarasjitt S. Spectral exponent characteristics of intracranial EEGs for epileptic seizure classification. IRBM. 2015;36(1):33–39. doi:10.1016/j.irbm.2014.07.005

64. Yu Z, Yang B, Wei P, et al. Critical biomarkers for responsive deep brain stimulation and responsive focal cortex stimulation in epilepsy field. Fundamental Research. 2025;5(1):103–114. doi:10.1016/j.fmre.2024.05.018

65. Kopf M, Martini J, Stier C, et al. Aperiodic Activity Indexes Neural Hyperexcitability in Generalized Epilepsy. eNeuro. 2024;11(9). doi:10.1523/ENEURO.0242-24.2024

66. Armstrong C, Zavez A, Mulcahey PJ, et al. Quantitative electroencephalographic analysis as a potential biomarker of response to treatment with cannabidiol. Epilepsy Res. 2022;185:106996. doi:10.1016/j.eplepsyres.2022.106996

67. Coa R, La Cava SM, Baldazzi G, et al. Estimated EEG functional connectivity and aperiodic component induced by vagal nerve stimulation in patients with drug-resistant epilepsy. Front Neurol. 2022;13:1030118. doi:10.3389/fneur.2022.1030118

68. Yang Y, Wang J, Wang X, et al. Long-term effects of vagus nerve stimulation on EEG aperiodic components in patients with drug-resistant epilepsy. Ther Adv Neurol Disord. 2024;17:17562864241279124. doi:10.1177/17562864241279124

69. Kundu B, Charlebois CM, Anderson DN, Peters A, Rolston JD. Chronic intracranial recordings after resection for epilepsy reveal a “running down” of epileptiform activity. Epilepsia. 2023;64(7):e135–e142. doi:10.1111/epi.17645

70. Marrosu F, Santoni F, Fà M, et al. Beta and gamma range EEG power-spectrum correlation with spiking discharges in DBA/2J mice absence model: role of GABA receptors. Epilepsia. 2006;47(3):489–494. doi:10.1111/j.1528-1167.2006.00456.x

71. Schroeder GM, Chowdhury FA, Cook MJ, et al. Multiple mechanisms shape the relationship between pathway and duration of focal seizures. Brain Commun. 2022;4(4):fcac173. doi:10.1093/braincomms/fcac173

72. Salami P, Borzello M, Kramer MA, Westover MB, Cash SS. Quantifying seizure termination patterns reveals limited pathways to seizure end. Neurobiol Dis. 2022;165:105645. doi:10.1016/j.nbd.2022.105645

73. Nakajima M, Halassa MM. Thalamic control of functional cortical connectivity. Curr Opin Neurobiol. 2017;44:127–131. doi:10.1016/j.conb.2017.04.001

